# Stress response in *Entamoeba histolytica* is associated with robust processing of tRNA to tRNA-halves

**DOI:** 10.1101/2021.09.30.462689

**Authors:** Manu Sharma, Hanbang Zhang, Gretchen Ehrenkaufer, Upinder Singh

## Abstract

tRNA-derived fragments have been reported in many different organisms and have diverse cellular roles such as regulating gene expression, inhibiting protein translation, silencing transposable elements and modulating cell proliferation. In particular tRNA halves, a class of tRNA fragments produced by the cleavage of tRNAs in the anti-codon loop, have been widely reported to accumulate under stress and regulate translation in cells. Here we report the presence of tRNA-derived fragments in *Entamoeba* with tRNA halves being the most abundant. We further established that tRNA halves accumulate in the parasites upon different stress stimuli such as oxidative stress, heat shock, and serum starvation. We also observed differential expression of tRNA halves during developmental changes of trophozoite to cyst conversion with various tRNA halves accumulating during early encystation. In contrast to other systems, the stress response does not appear to be mediated by a few specific tRNA halves as multiple tRNAs appear to be processed during the various stresses. Furthermore, we identified some tRNA-derived fragments are associated with *Entamoeba* Argonaute proteins, *Eh*Ago2-2, and *Eh*Ago2-3, which have a preference for different tRNA-derived fragment species. Finally, we show that tRNA halves are packaged inside extracellular vesicles secreted by amoeba. The ubiquitous presence of tRNA-derived fragments, their association with the Argonaute proteins, and the accumulation of tRNA halves during multiple different stresses including encystation suggest a nuanced level of gene expression regulation mediated by different tRNA-derived fragments in *Entamoeba*.

**Importance:** tRNA-derived fragments are small RNAs formed by the cleavage of tRNAs at specific positions. These have been reported in many organisms to modulate gene expression and thus regulate various cell functions. In the present study, we report for the first time the presence of tRNA-derived fragments in *Entamoeba*. tRNA-derived fragments were identified by bioinformatics analyses of small RNA sequencing datasets from the parasites and also confirmed experimentally. We found that tRNA halves accumulated in parasites exposed to environmental stress or during developmental process of encystation. We also found that shorter tRNA-derived fragments are bound to *Entamoeba* Argonaute proteins, indicating that they may have a potential role in the Argonaute-mediated RNA-interference pathway which mediates robust gene silencing in *Entamoeba*. Our results suggest that tRNA-derived fragments in *Entamoeba* have a possible role in regulating gene expression during environmental stress.

## Introduction

*Entamoeba histolytica* is a protozoan parasite with a biphasic lifecycle. A dormant cyst stage causes infection upon ingestion of contaminated food or water. The cysts transform into invasive trophozoites in the small intestine and proliferate upon reaching the colon (1). *Entamoeba* parasites can elicit disease symptoms if trophozoites invade the colonic wall, causing colitis. The life cycle is completed when trophozoites convert to cysts in the colon through a process known as encystation; these newly formed cysts can be excreted and spread to new hosts. Currently, the signals that cause the trophozoites to encyst in the colon are not completely understood (2). Our group has previously shown that extracellular vesicles (EVs) secreted by encysting parasites can promote encystation in *Entamoeba* parasites, though it was not ascertained what factors in the EV cargo were responsible for modulating encystation (3).

tRNA genes are extremely abundant in the amoeba genome with approximately 4,500 copies. The tRNA genes are organized in unique tandem repeats clustered into arrays that make up over 10% of the genome. Though the function of the arrays is not yet completely clear, there is evidence to show that these have a structural role in amoeba in the absence of classical telomeres (4, 5). Only four tRNA genes are outside of the arrays and the rest are exclusively found in the arrays. The existence of these low copy number tRNA genes does not affect efficient translation and no correlation has been found between codon usage and tRNA copy number.

tRNA-derived fragments (tRFs) are a class of small RNAs, approximately 16-40 bases in length, that have been identified in organisms from all domains of life (6). tRFs are produced through the cleavage of either precursor or mature tRNAs. Studies have shown that the cleavage can occur at any position on the tRNAs and is “context-dependent” (7). The best characterized tRFs are the tRNA halves, formed by the cleavage at the anticodon loop of mature tRNAs (8, 9). tRNA halves are 30 to 40 nucleotides in size and map to either the 5’ or 3’ end of the mature tRNA. In humans, angiogenin can cleave tRNAs at the anticodon loop during stress; the resultant tRNA halves have been shown to regulate translation (10). However, angiogenin independent mechanisms of tRNA halves have also been reported in human cells, as well as in other organisms that lack this protein (11, 12). Other tRFs are tRF-3s, tRF-5s, tRF-1s, and misc-tRFs (6). misc-tRFs include the internal-tRFs (itRFs) that span the internal regions of mature tRNAs, but do not map to either the 5’ or 3’ ends. tRFs generated from the 3’-end of mature tRNAs contain a 3’-CCA tail that is added to the precursor tRNA upon maturation.

One of the first studies in parasites describing tRNA fragments was performed in *Giardia lamblia* (13). Li et al. described a class of small RNAs, approximately 46nt in length, derived from the 3’ end of mature tRNAs. These small RNAs, referred to as stress induced tRNAs (sitRNAs), contained a 3’-CCA tail and accumulated in the cell during encystation. Si-tRNAs are longer than tRNA halves and contain a cleavage site in the anticodon right arm. It was observed that si-tRNAs were generated indiscriminately from the entire tRNA family in the parasite. The authors further showed that si-tRNAs also accumulated during various stress-stimuli such as temperature shock and serum deprivation. More recently, tRNA halves have been shown to modulate translation during stress response in *Trypanosoma brucei* (12). Fricker et al. demonstrated that during serum starvation, 3’ tRNA^Thr^ accumulated in parasites and associated with ribosomes and polysomes to stimulate global protein translation (12). tRFs have also been reported in other parasites such as *Trypanosoma cruzi* and *Plasmodium falciparum* (14, 15). Moreover, *Leishmania* parasites and *T. cruzi* are known to package tRFs in their secreted exosomes suggesting that these parasites could be using tRFs as means of intercellular communication (16, 17).

tRFs have also been shown to associate with Argonaute (Ago) proteins in multiple organisms (18–21). This association suggests a possible role of tRFs in the RNA interference (RNAi) pathway. Indeed, there is increasing evidence demonstrating the loading of tRFs onto Argonaute proteins resulting in transcript cleavage based on sequence complementarity (22, 23). In humans, the 3′-tRF from tRNA^GlyGCC^ has been shown to bind with Ago and lead to degradation of RPA1 to arrest B cell lymphoma (24). In the *E. histolytica* genome, three Ago proteins have been identified (EHI_125650, EHI_186850, and EHI_177170); *Eh*Ago2-2 (EHI_125650) is the most highly expressed Ago protein in *E. histolytica*. We have previously demonstrated *Eh*Ago2-2 binds to small RNA populations of 27nt or 31nt in size and mediates transcriptional gene silencing (25, 26).

In our present work, we wished to identify if tRFs are present in *Entamoeba* and if there is a change in abundance in response to stress. We determined that tRFs can be identified in small RNA sequencing datasets of *Entamoeba*, and that these confirm experimentally by Northern blot experiments. Additionally, we determined that tRNA halves accumulate in amoeba during stress and encystation. Furthermore, we observed short tRFs (24-30nt in size) bound to Eh Argonaute proteins, suggesting a possible role in the Argonaute mediated gene silencing pathway. However, unlike in some other systems, such as *Trypanosoma cruzi*, where the stress response is mediated by elevation of only a few (or predominantly one) tRNA halves, stress conditions in amoeba leads to the elevation of a large subset of tRNA halves suggesting that the tRNA fragment pool in this parasite is more diverse and varied and possibly working in a redundant manner to carry out any molecular regulatory role after stress induction.

## Results

### tRNA fragments identified in *E. histolytica* small RNA libraries and confirmed by Northern blot analyses

To check for the presence of tRNA fragments in *Entamoeba*, we interrogated a previously published small RNA-seq datasets. These datasets consisted of small RNA libraries generated from *E. histolytica* parasites under basal growth conditions and during two different stresses (oxidative stress or heat stress (27, 28)). For the purpose of our previous work, the libraries had been prepared by fractionating the total RNA into two size fractions – 15-30nt and 30-45nt. In the present study, the sequencing datasets from the two size-selected groups (for each sample) were combined together for subsequent analysis. For mapping the sequence reads, an index of *E. histolytica* tRNA sequences was first created from genomic sequences using tRNAscan-SE (29). Mature tRNAs contain a non-templated “CCA” tail at the 3’ end. As such, tRNA fragments derived from the 3’ ends of mature tRNAs contain a 3’ CCA tail, whereas those derived from immature pre-tRNAs do not. A ‘CCA’ tail was added to the 3’ terminal end of these predicted tRNA sequences to create a final tRNA index. The tRNA sequence index was used for mapping the sequencing datasets mentioned above, using Bowtie version 1.0.0 using a previously established pipeline (27, 30).

The total number of reads for each sample and the percentage of reads mapping to tRNAs is shown in Fig. 1A. Similar-sized libraries were obtained for the control and oxidative stress treated samples (1,028,524 and 1,446,363, respectively), and much higher for the heat-shock samples (2,388,635). As published earlier, almost 40% of the reads were unique reads indicating that the sequencing was not saturated and likely missing rare species (27). Overall, around 3-6% of the total sequence reads mapped to the tRNAs. The sequence reads that mapped to the tRNA sequences were selected and analyzed based on their lengths (Fig. 1B). A non-random distribution was observed, suggesting that the tRNA fragments were not generated by indiscriminate degradation. The peaks ∼35nts corresponded to the expected size for tRNA halves. Other tRFs, besides tRNA halves, were also present in the sequence reads as seen in peaks around 26 and 29nts. A majority of these tRNA fragments mapped to the 5’ end of their respective tRNAs (> 90% for each dataset). This trend is not uncommon, and for example it was recently reported that tRFs originating from the 5’ end constituted around 86% of total tRFs in *Plasmodium falciparum* (14). However, there could have been a further bias generated during our library preparations which involved a 5’ phosphate-dependent cloning step. As a result of this, tRFs containing 5’ phosphate groups would be preferentially cloned compared to those formed by cleavage of tRNA resulting in a 5’-OH group.

**Figure 1:**
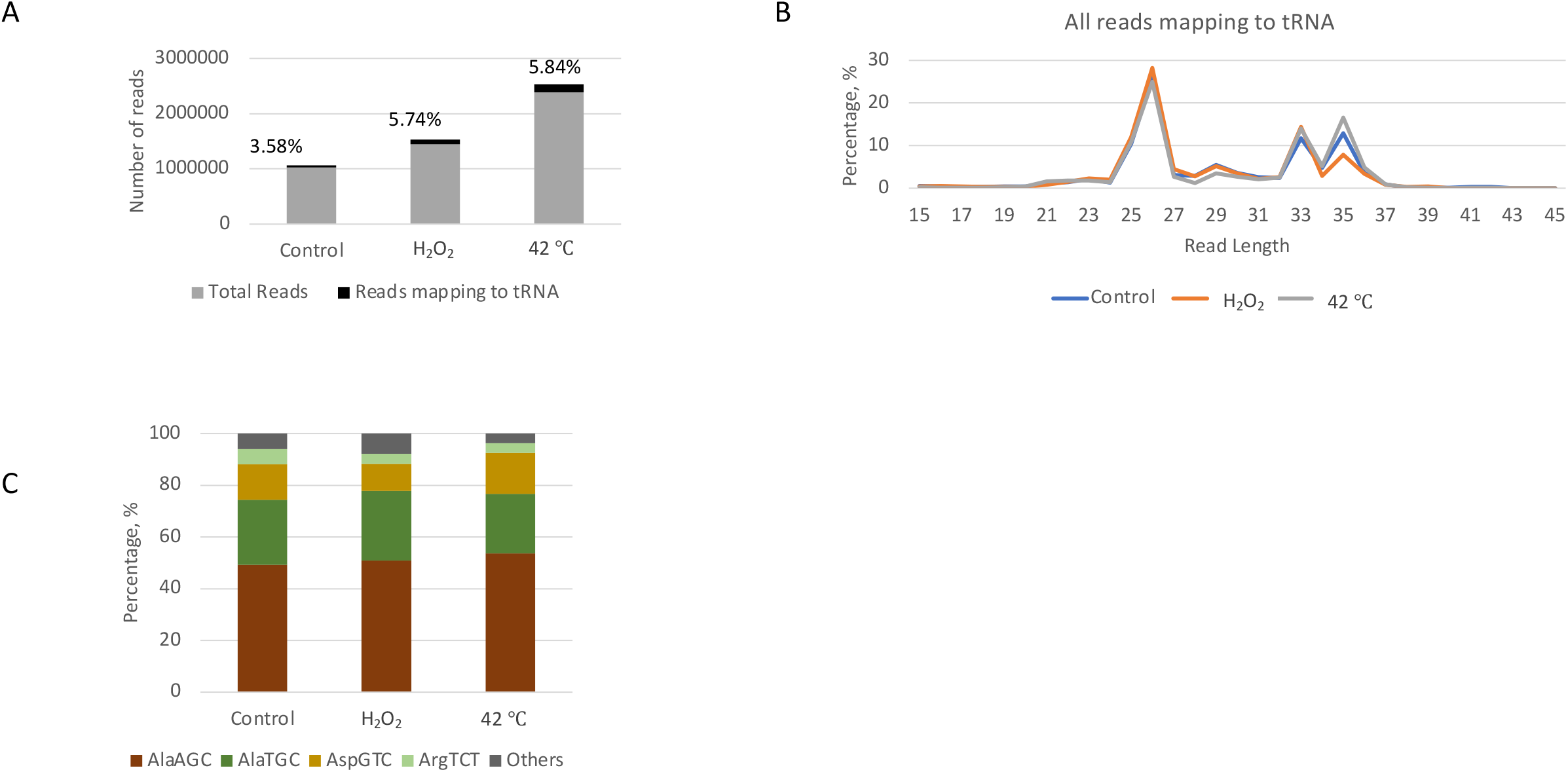
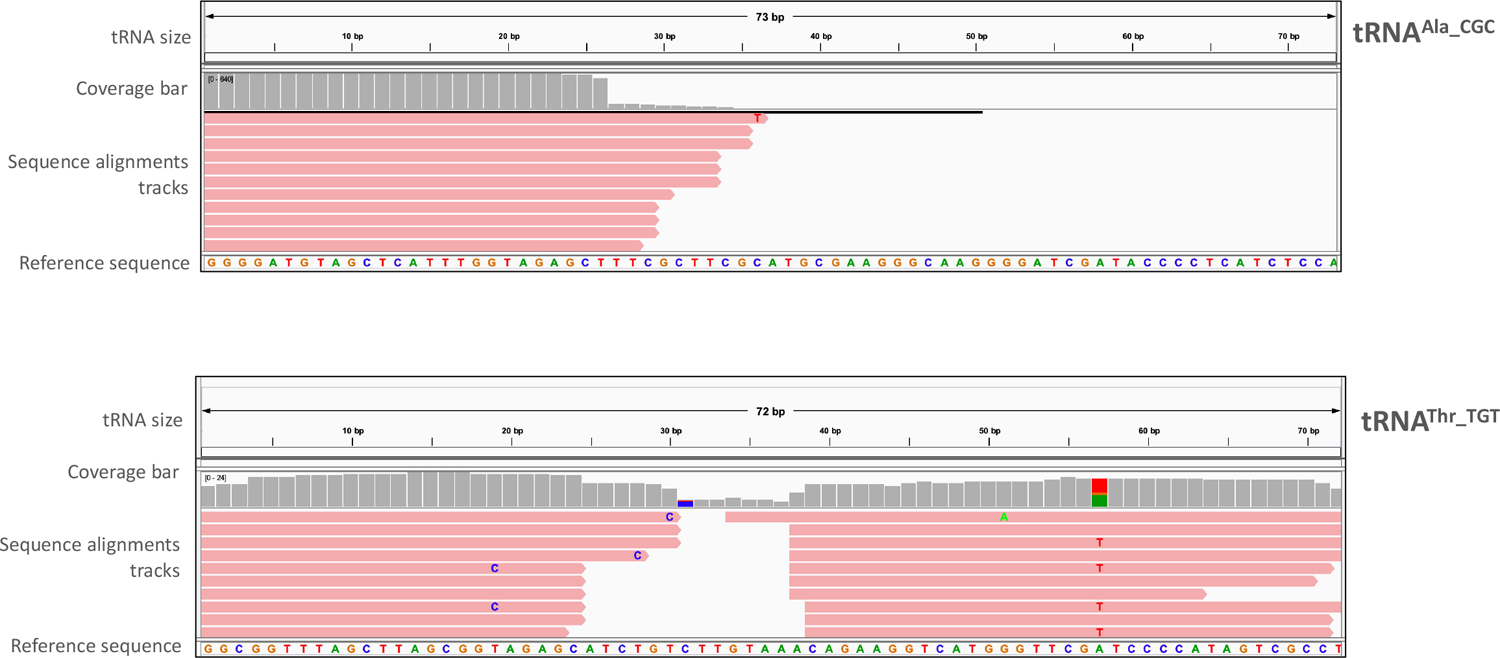
Bioinformatics analyses of small RNA sequencing dataset from *E. histolytica* demonstrates the presence of tRFs. **(A)** Small RNA sequences generated from size selected libraries from *E. histolytica* in either basal or stress conditions were mapped to tRNA sequences using Bowtie. Chart showing the percentage of sequence reads that map to tRNA in different samples. **(B)** Length distribution of the different reads that mapped to tRNA sequences. **(C)** Frequency of reads mapping to the four most abundant tRNAs seen. **(D)** Representatitive IGV mapping of sequence reads from *E. histolytica* (oxidative stress sample) mapping against tRNA templates-tRNA^Ala_CGC^ and tRNA^Thr_TGT^. The top of each IGV mapping shows the size of the parent tRNA template (72 or 73 nt for tRNA^Ala_CGC^ and tRNA^Thr_TGT^ respectively). The coverage bar in grey displays the depth of the reads at each locus as a bar chart. Below this, the sequence alignment tracks represent each sequence read aligned to the parent tRNA, whose sequence is shown as the “Reference Sequence” below. As seen for a majority of tRNAs, both 5’ and 3’ tRNA halves were observed for tRNA^Thr_TGT^. However, only the 5’ tRNA half was observed for tRNA^Ala_CGC^.

The frequency of the tRNAs to which the reads mapped (note that these data show the cumulative mapping for all sequence reads of different sizes) is in Fig. 1C. The tRNA frequency is identical for the control sample compared to the samples under different stresses. The tRFs derived from 4 different tRNAs (tRNA^Ala_AGC^, tRNA^Ala_TGC^, tRNA^Asp_GTC^, and tRNA^Arg_TCT^) constituted around 80% of the total reads and tRNA^Ala_AGC^ was the most abundant tRNA to which the sequence reads mapped. Closer inspection of the sequence reads showed that the data are skewed by multiple copies of a few reads, similar to what has been reported for other systems.

Approximately 10% of the sequence reads mapped to the 3’ end of tRNAs. We found that approximately 80% of these reads that mapped to the 3’ end also contained a CCA tail, suggesting the tRFs were formed predominantly by the cleavage of mature tRNAs. The alignment of the sequence reads to representative tRNAs (tRNA^Ala_CGC^ and tRNA^Thr_TGT^,) in the integrated genome viewer (IGV) is in Fig. 1D (31). We are not comparing the tRFs from different samples but as a means to display the different tRNA-derived fragments in our dataset. Therefore, only the sequence reads from the oxidative-stress sample are used to show the mapping of different tRFs with respect to the parent tRNA. As seen in Fig 1D, the mapping is not randomly distributed but instead is localized to specific regions on the parent tRNA template (predominantly around the 5’ end) which suggests that these reads are not degradation products. However, the diversity in the positions of termini suggests that some of the reads maybe the results of degradation/exonucleolytic process.

To confirm the presence of the tRFs in *E. histolytica* and to compare the tRF levels during different stress stimuli, northern blot analysis of total RNA was performed. Northern blot analysis for 5’ tRNAF^Ala_CGC^ and 5’tRNAF^ARG_ACG^, displaying full length tRNA, and tRFs of different sizes are shown (Fig. 2A). tRNA halves (around 30-35nt in size) were the most abundant tRFs and were found to accumulate under the different stress conditions tested. Distinct bands corresponding to smaller tRFs were also observed but their relative amounts appeared to remain unaffected during the stress as compared to the control. We checked for tRNA halves corresponding to other tRNAs and found that in each case there was a stress-induced accumulation of tRNA halves (Fig. 2B). Oxidative stress following H_2_O_2_ treatment had the most significant impact on the accumulation of different tRNA halves. tRNA halves were not observed for 3’ tRNA^Ala_CGC^. The presence of 5’ tRNA^ALA^ portion but the absence of its corresponding 3’ half has also been observed in other systems (32).

**Figure 2:**
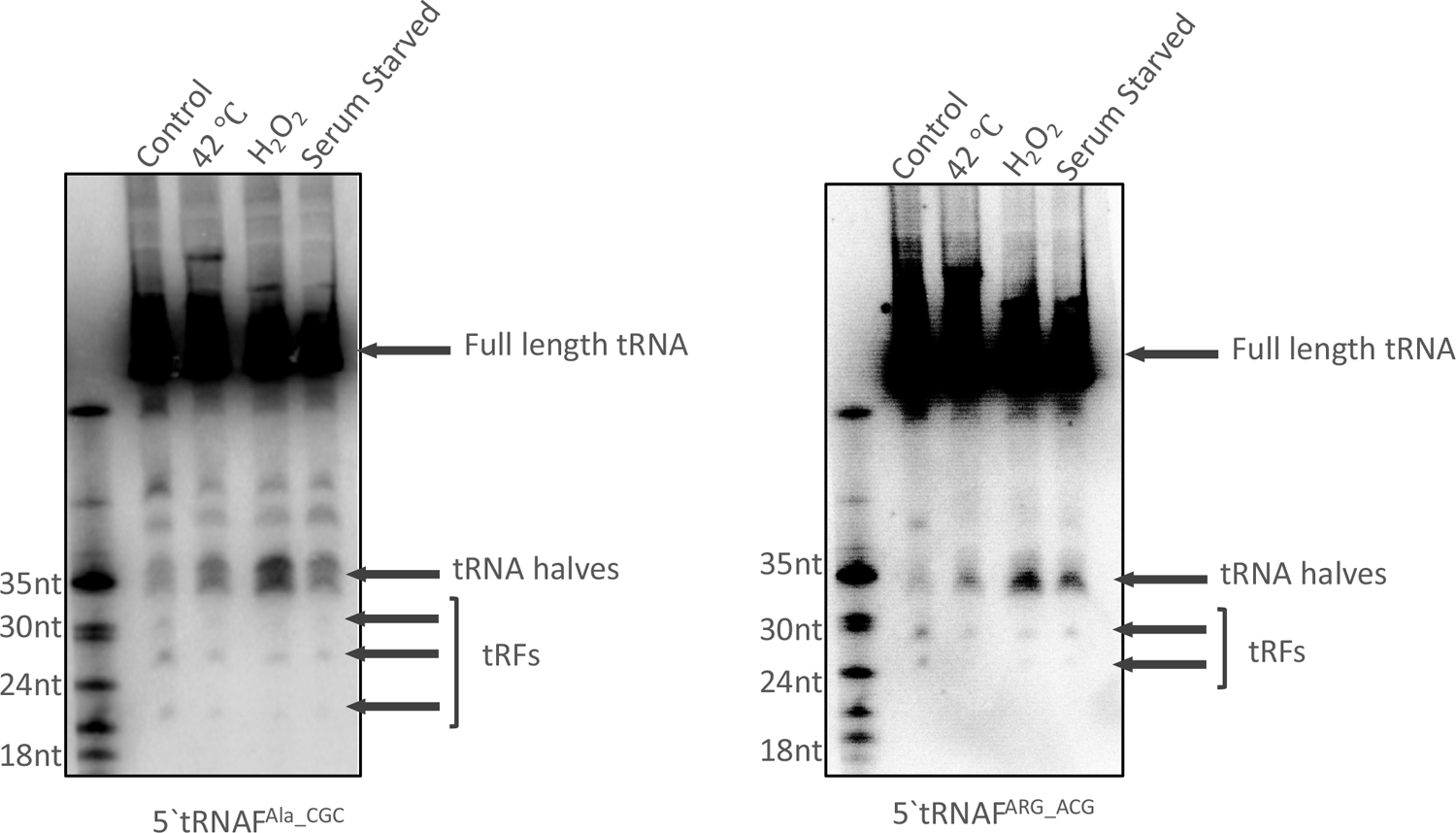
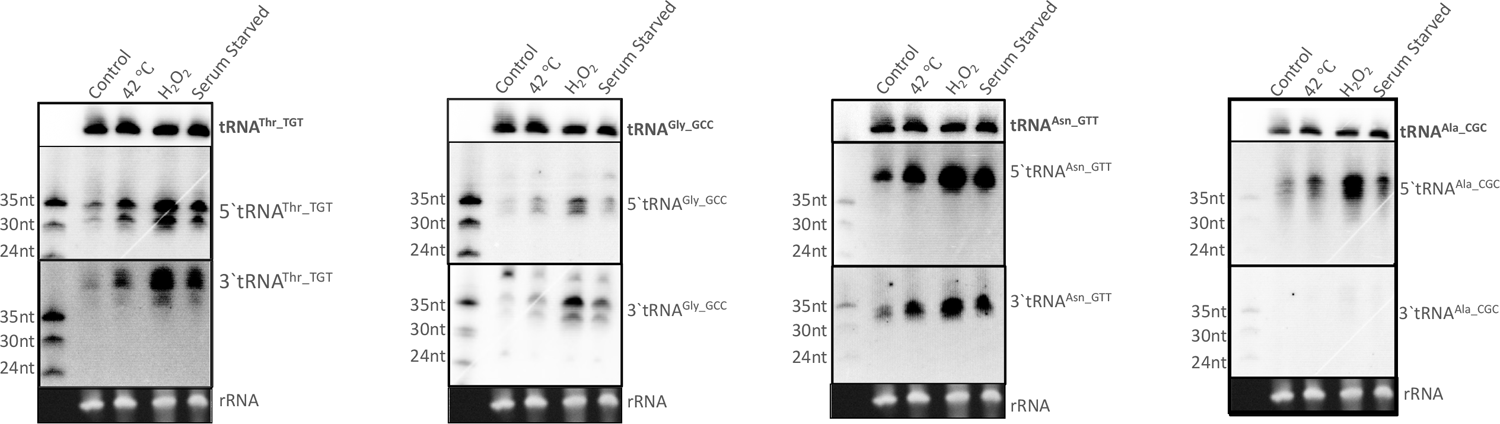
Northern blot analysis of tRNA fragments from *E. histolytica* shows accumulation of tRNA halves during stress. **(A)** Total RNA was prepared from *E. histolytica* parasites under basal condition or stressed with oxidative stress (H_2_O_2_), Heat shock (42 C) or serum starvation, and analyzed by northern blotting. Representative blots of 5’tRNAF^Ala_CGC^ and 5’tRNAF^ARG_ACG^ show the presence of abundant tRNA halves during stress as well as smaller tRFs of distinct sizes. **(B)** Northern blots showing tRNA halves for different tRNAs. Both 5’ and 3’ tRNA halves were seen to accumulate in response to the different stresses, for tRNA^Thr_TGT^, tRNAF^Gly_GCC^, tRNAF^Asn_GTT^ whereas only the 5’ tRNA half was observed for tRNAF^Ala_CGC^.

### tRNA halves accumulate in *Entamoeba* parasites during encystation

Since *Entamoeba* encystation also involves stress on the parasites in the form of nutritional depletion, we wanted to determine if tRNA halves are regulated during this developmental process. Efficient encystation in *E. histolytica* has not been achieved *in vitro* and encystation of *Entamoeba* is studied using *E. invadens* (33, 34). We performed bioinformatics analyses of small RNA sequencing datasets (size selected RNA < 45nt in length) comparing developmental stages (trophozoites, early and late cysts, and excysting parasites) in *E. invadens* (27, 35). We observed that ∼1 – 4% of sequence reads mapped to the tRNA sequences (Fig. 3A). Analysis of the length distribution showed that tRFs were present in all the different developmental stages with peaks at around 25nt and 34nts (Fig. 3B). tRNA halves comprised a larger fraction of the total reads mapping to tRNAs in early cysts and excysting parasites compared to trophozoites. The 25nt sequence reads were the most abundant tRNA fragment species in trophozoites. As observed with *E. histolytica* parasites, a majority of the reads mapped to the 5’ region of tRNAs (>80%). The frequency of reads mapping to different tRNAs was found to be similar across different stages, with a few tRNAs (tRNA^Asp_GTC^, tRNA^Gln_CTG^, tRNA^Gln_TTG^, and tRNA^Cys_GCA^) accounting for a majority of the sequence read-mapping (Fig. 3C). However, there was some apparent variability in the tRFs belonging to different stages of development. For example, reads mapping to tRNA^Asp_GTC^ constituted around 20% of total reads for trophozoites, early and late cysts but were only 5% of the reads in excysting parasites.

**Figure 3:**
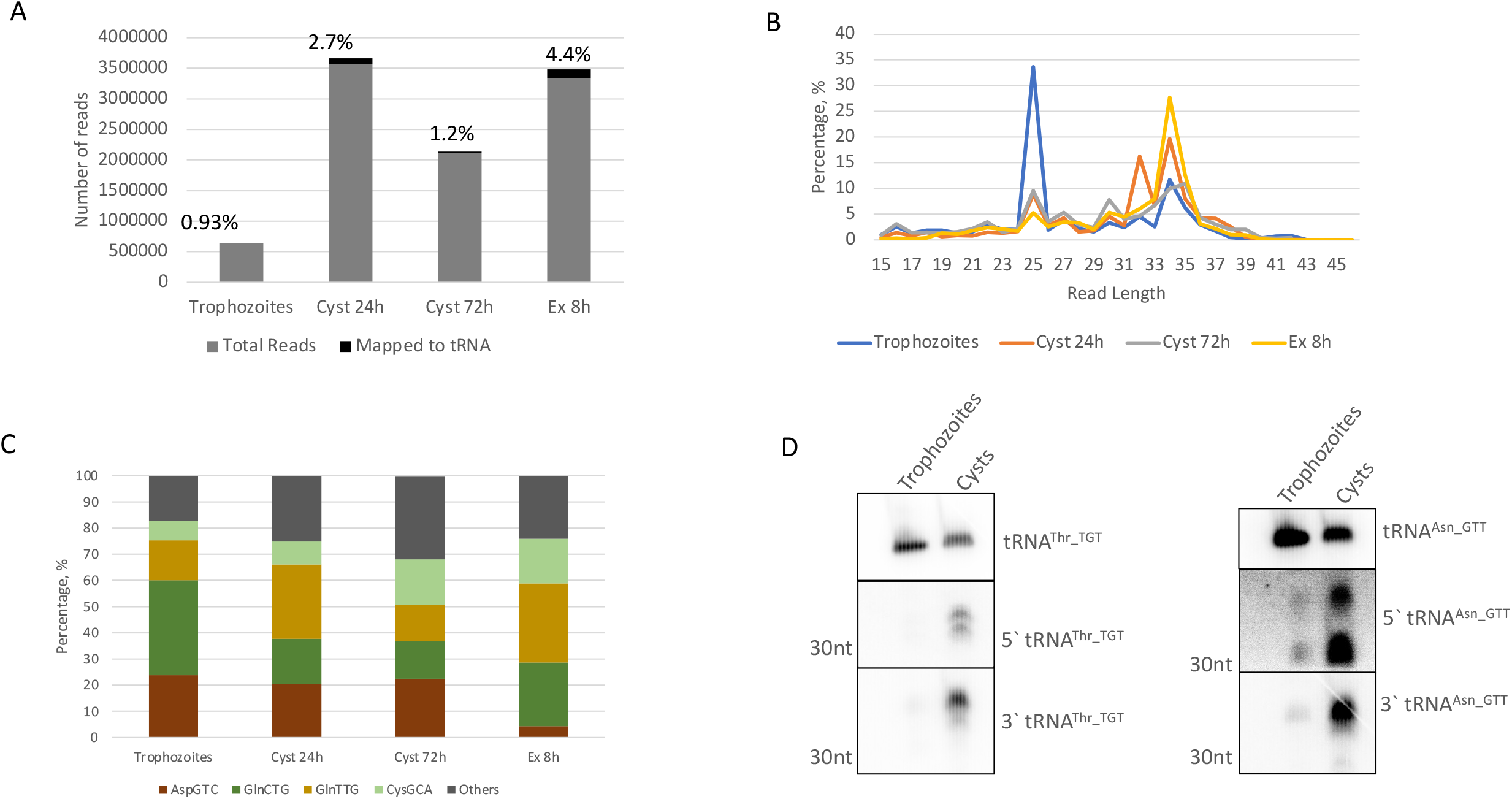
tRNA halves accumulate during encystation of *E. invadens*. **(A)** Small RNA sequences generated from size selected libraries from *E. invadens* parasites in different stages-Trophozoites, 24h cysts, 72 h cysts or excysting parasites at 8h. Chart shows percentage of reads mapping to tRNA. **(B)** Length distribution of reads mapping to tRNA sequences. **(C)** Frequency of reads mapping to most abundant tRNAs. **(D)** Northern blot analysis of total RNA from 24 h cysts and trophozoites probed for tRNA halves. Both 5’ and 3’ halves were seen to be accumulated in the early cysts stage compared to the trophozoites

Northern blot analysis was carried out to compare the tRNA halves in *E. invadens* trophozoites and early cysts (24 hrs following the induction of encystation). Similar to the stress response in *E. histolytica*, various tRNA halves accumulated during encystation. Representative northern blots for the 5’ and 3’ halves of tRNA^Thr_TGT^ and tRNA^Asn_GTT^ are shown (Fig. 3D). Northern blots carried out for 5’ tRNA halves of tRNA^Arg_TCT^, tRNA^Ala_CGC^, tRNA^Gly_GCC^, and tRNA^Ala_AGC^ are shown in Supplementary Fig S1.

### tRNA fragments in extracellular vesicles derived from *E. histolytica*

In our previous work, we characterized EVs prepared from amoeba and analyzed their RNA cargo by sequencing (3). In the present study, the RNA-seq dataset from the size selected EV-RNA library was analyzed for tRFs as described above. Around 1% of the sequence reads mapped to tRNAs (Fig. 4A). Similar to what was observed in Fig. 1B, the length distribution of the cellular RNAs had peaks at around 26, 29, and 33nts Fig. 4B. The EV RNA had peaks at 29 and a more pronounced peak at 33nts, suggesting that tRNA halves are the most abundant tRNA derived fragments packaged in these EVs. Note that the protocol for EV preparation required the use of serum free media and therefore, both the EV and cellular RNA samples had been serum starved (3). As seen in Fig. 1D, serum starvation is one of the stressors that lead to accumulation of tRNA halves, similar to what was reported in different systems (12, 36). The frequency of the reads mapping to different tRNAs is in Fig. 4C. As previously seen, a majority of the reads mapped to a few tRNAs (tRNA^Ala_AGC^, tRNA^Ala_TGC^, tRNA^Arg_TCT^, tRNA^Asp_GTC^). We did not find any major differences in the IGV profiles for EV and cellular reads for the different tRNAs (data not shown). Northern blot analysis was performed to check for tRNA halves in *E. histolytica* cellular and EV RNA. EV preparation was done in serum free conditions to exclude contamination from exogenous exosomes in the serum (3). RNA isolated from the EVs was compared to cellular RNA isolated from parasites grown under similar serum free conditions. As seen in Fig. 4D, the 3’tRNA^Asn_GTT^ tRNA half was observed in the EV RNA.

**Figure 4:**
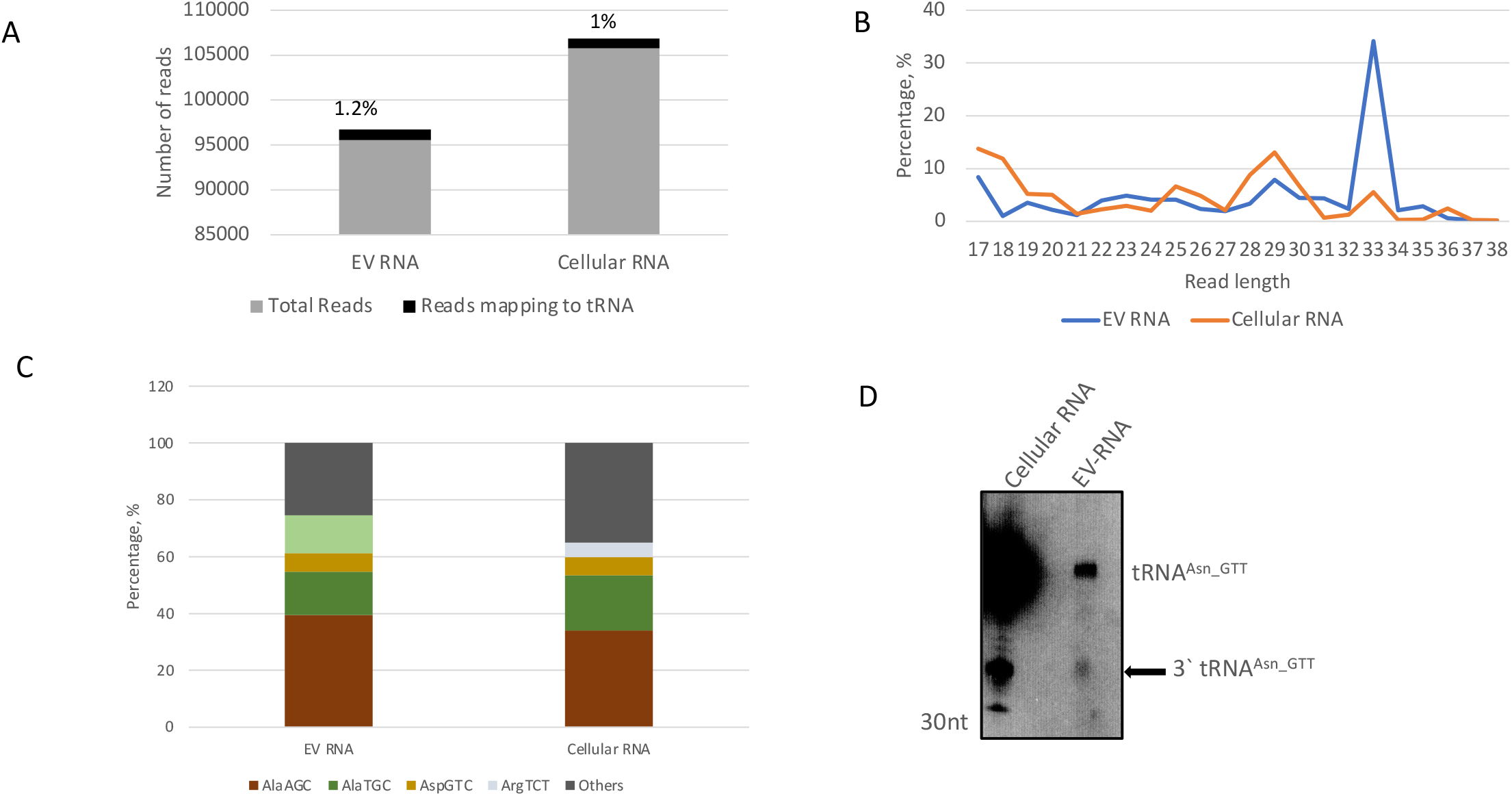
tRNA fragments are identified in small RNA sequence datasets obtained from size selected libraries of EVs secreted by *E. histolytica*. **(A)** Percentage of reads mapping to tRNA. **(B)** Length distribution **(C)** RNA frequency. **(D)** Northern blot analysis of EV and cellular RNA prepared from *E. histolytica* parasites. 3’tRNA^Asn_GTT^was seen to be packaged in EV.

### tRNA-derived fragments are associated with *E. histolytica* Argonaute (EhAgo) proteins

We previously demonstrated that two small RNA populations, 27 and 31nt in size, associate with *Eh*Ago proteins (25, 26, 28). Since Ago proteins have been reported to bind tRNA fragments in other systems (23, 37), we analyzed our sequencing data from size-fractionated total RNAs isolated from IPs of each of the 3 *Eh*Agos: Ago2-1, Ago2-2 and Ago2-3. We found that the number of reads mapping to tRNA in IP-RNA was comparable to what we observed in whole cell sRNA samples. However, there were relatively higher mapping of reads from Ago2-3 (5.8%) compared to those from Ago2-2 (3.8%) or Ago 2-1 (1%) (Fig. 5A). Length distribution of the mapped reads showed a non-random distribution with peaks at around 24 and 27nt for these tRNA fragments Fig. 5B. A majority of the sequence reads from the Ago2-3 samples were 24nt in length, whereas Ago2-2 and Ago2-1 were predominantly 27nt in size. Larger tRFs (e.g. tRNA halves around 33nt) were not observed binding to the Ago suggesting that Ago proteins might have a role in guiding shorter tRNA-derived fragments; this phenomena has been reported in other systems (37). The mapped reads were viewed on IGV against tRNA templates and two representative tRNAs-tRNA^Ala_TGC^ and tRNA^Arg_TCT^ are shown in Supplementary Fig. S2. The 5’ tRNA-derived fragments were the most abundant group of tRFs binding to the three Ago proteins. Interestingly, Ago2-2 seems to bind to itRF species for different tRNAs. Though we did not observe itRFs in the cellular small RNA dataset (analyzed in Fig. 1), it is possible that the itRFs were enriched and stabilized in the Ago overexpression cell lines that were used for the IP experiments.

**Figure 5:**
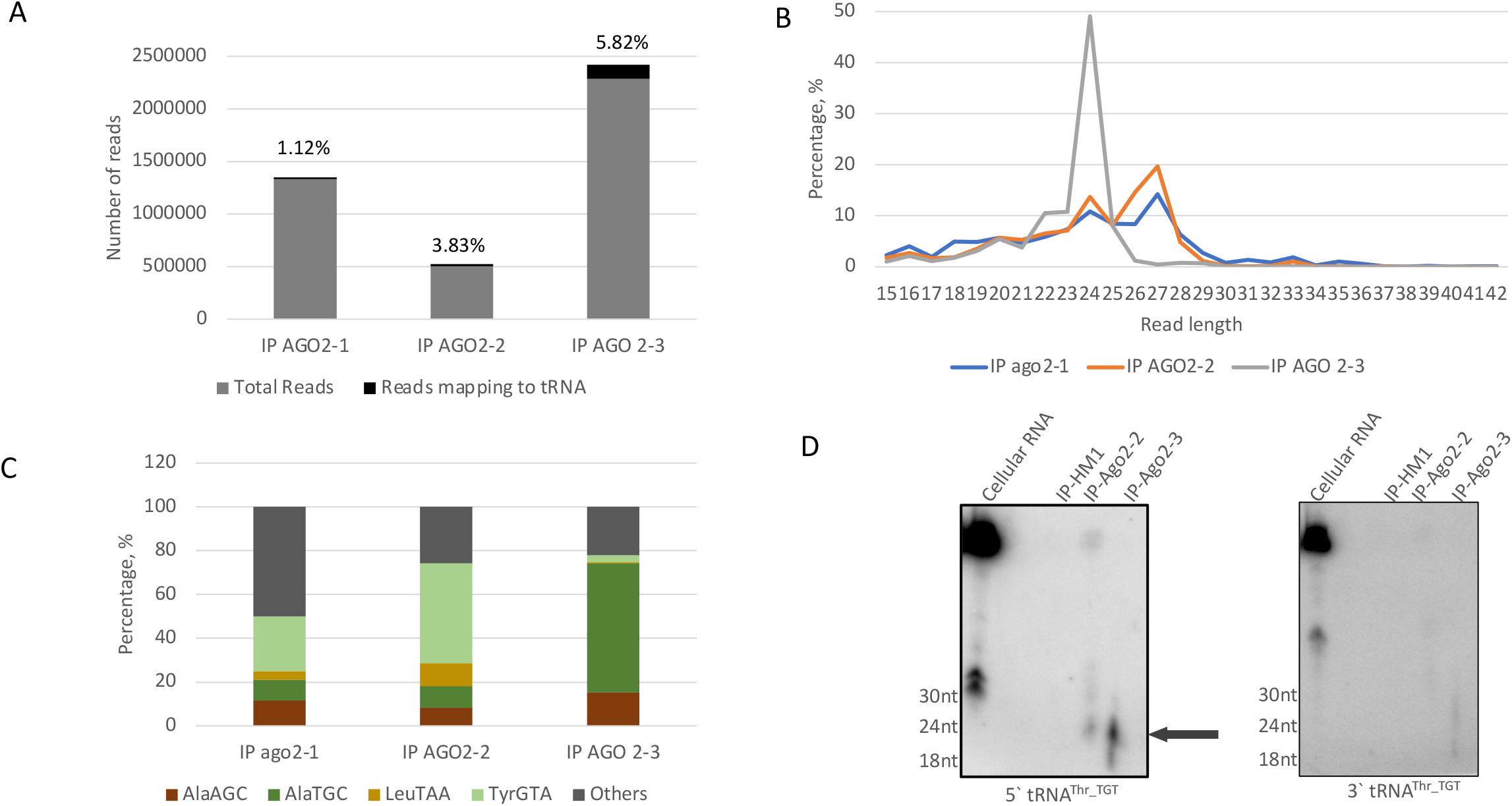
tRFs associate with the three *Eh*Ago proteins in amoeba. **(A)** IP from Myc-tagged overexpression cell lines for the three *Eh*Ago proteins were performed and RNA from beads isolated. Small RNA libraries were prepared by size selection and sequenced. Chart showing the percentage of sequence reads that map to tRNA in different samples. **(B)** Length distribution of the different reads that mapped to tRNA sequences. **(C)** Frequency of reads mapping to the most abundant tRNAs. **(D)** Norther blot analysis of IP-RNA prepared from myc-*Eh*Ago2-2 and myc-*Eh-*Ago2-3 overexpression cells lines. IP-HM1 refers to IP performed with un-transfected HM1 control cells. Cellular RNA was used as a positive control. tRFs were observed in both IP-Ago2-2 and IP-Ago2-3 samples for 5’ tRNA^Thr_TGT^ but not for 3’ tRNA^Thr_TGT^.

Northern blot analysis was carried out for the Ago-IP samples. We focused on association of tRFs with *Eh*Ago2-2 & *Eh*Ago2-3 since those had the largest populations of mapping sequence reads. *E. histolytica* cell lines overexpressing Myc-tagged *Eh*Ago2-2 or *Eh*Ago2-3 were used for anti-Myc immunoprecipitation and the total associated RNA was isolated as described (28). Anti-Myc IP was carried out in parallel on untransfected *E. histolytica* trophozoites and RNA isolation performed to obtain a negative control. 5’ tRNA^Thr_TGT^ -derived fragments, around 27nt in length, were observed in RNA isolated from both *Eh*Ago2-2 and *Eh*Ago2-3 IPs, but with a stronger association with the latter (Fig. 5D). Compared to 5’-tRNA^Thr_TGT^-derived fragments, accumulation of other tRNA-derived fragments was seen at a much lower intensity not perceptible in our northern blots. We could not identify tRNA-derived fragments from 3’-tRNA^Thr_TGT^. This would make sense if binding of small RNAs to Ago proteins is dependent on the 5’phosphate groups on RNAs.

In mammalian cells, generation of tRNA halves have been reported by angiogenin-dependent and angiogenin-independent mechanisms (11). However, we could not find homologs of angiogenin in the amoebic genome based on sequence similarity by reciprocal blast searches or presence of a conserved domain by the use of Pfam (data not shown). This could suggest that an alternative tRNA cleavage mechanism is used to generate the tRNA halves in *Entamoeba* parasites. Mechanisms for tRNA generation is not uncommon and have been reported in various organisms. In yeasts, the RNase T2 family member, Rny1, can cleave tRNAs at the anticodon loop, to generate tRNA halves (38). In *T. brucei*, no known homologs of angiogenin or Rny1 exist (12). *Entamoeba* species have RNases that belong to the RNase T2 family and it is possible that these proteins could be involved in tRNA half biogenesis in response to stress. Further work would be required to investigate this.

A schematic of our findings of tRFs in amoeba is shown in Fig. 6. Cleavage of tRNAs gives rise to tRFs of distinct sizes that appear to be precisely generated and are not degradation products. When the parasites are treated to different stress inducers, tRNA halves are the most abundant tRFs that accumulate. Both 5’ and 3’ tRNA halves were generated in response to stress, and these corresponded to various tRNA isoacceptors. Moreover, tRNA halves are packaged inside extracellular vesicles where they could have a potential role in extracellular communication with other parasites. As reported in other systems, tRFs could bind to Ago proteins to initiate gene silencing in a manner similar to microRNAs. The role of the smaller tRFs in amebic biology is not clear although we found that some of the smaller tRFs associate with the *Eh*Ago proteins.

**Figure 6:**
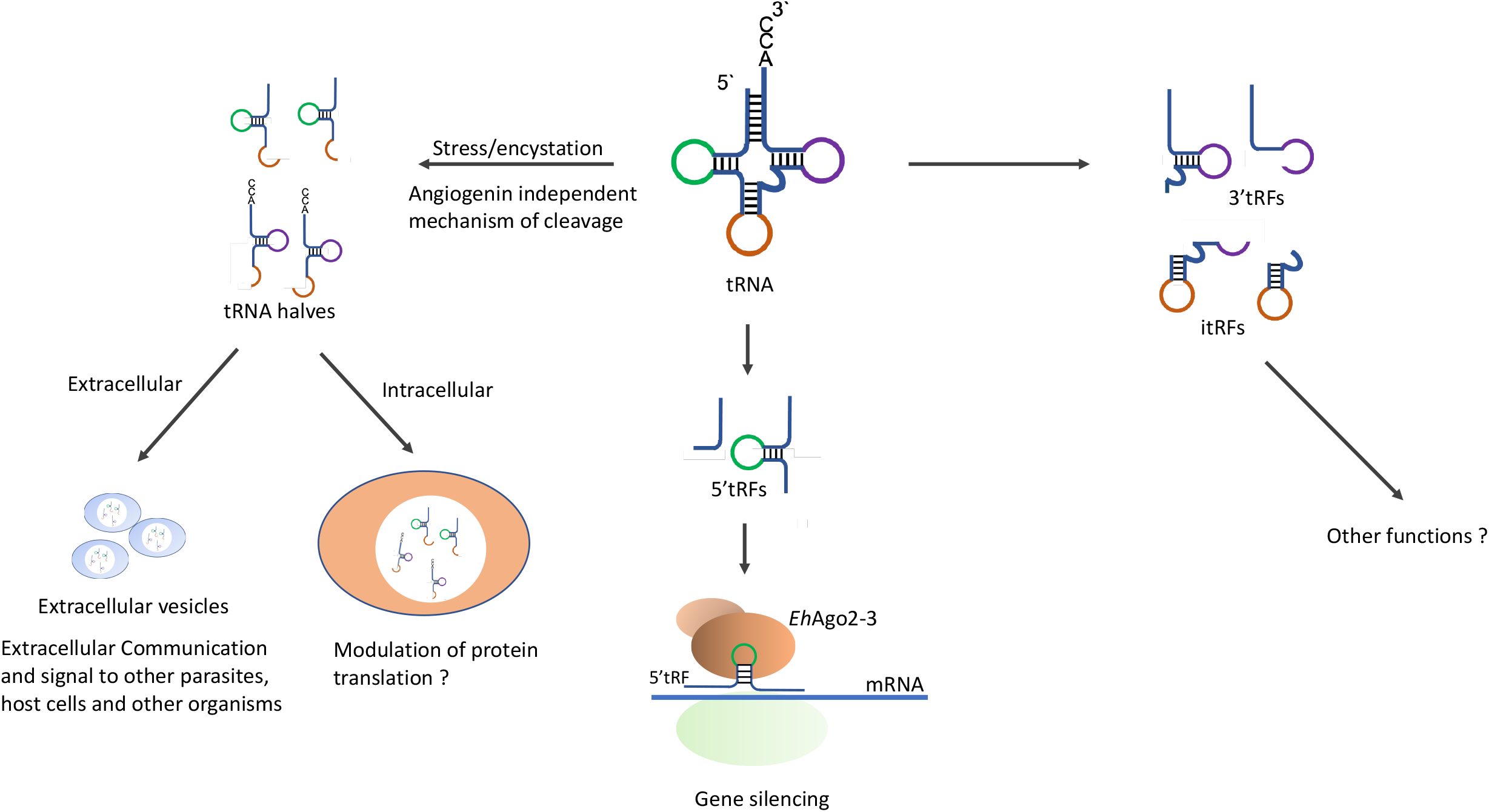
Schematic outlining potential roles of tRFs in amoeba. Both 5’ and 3’ tRFs are produced in amoeba. tRNA halves get accumulated during stress and during encystation and these can be subsequently packed into EVs for extracellular communication. Smaller tRFs – 3’, 5’ or itRFs are also present in Amoeba and are seen to bind to Ago proteins for possible roles in gene silencing.

## Discussion

tRNA genes are amongst the oldest genes that can be traced back to the putative RNA world that predated the separation of the three domains of life (39). It is no surprise then that tRNA-derived fragments have now been reported in Bacteria, Archaea, and Eukarya. Different classes of tRFs formed from the cleavage of either the mature or precursor tRNAs exist and have been shown to regulate gene expression in diverse contexts. Our present work provides evidence that multiple tRF species exist in amoeba and may have a potential to regulate gene regulation.

The amoeba genome is made up of arrays containing tandem repeats of tRNA clusters that make up over 105 of the genome (4). In spite of the high copy number of the tRNA genes, small RNA sequencing datasets in our labs have shown only around 5% of reads mapping to the tRNA (35, 40). Around 80% of the tRFs originated from just four tRNAs-tRNA^Ala_AGC^, tRNA^Ala_TGC^, tRNA^Asp_GTC^, and tRNA^Arg_TCT^. This has been reported in other systems as well (41–43). However, the tRF abundance did not correlate with either the codon usage or the tRNA copy numbers. Again, this is not surprising and other groups have reported a similar absence of correlation between codon usage and tRF abundance (44, 45). Ghosh et.al. had carried out a comprehensive analysis of available *E. histolytica* genome datasets to determine codon bias and found that the genome is AT rich and overall A or T ending codons are strongly biased in the coding region. However, when only considering highly expressed genes, there was a clear bias for C ending codons, suggesting that these codons are translationally optimal for amoeba (46). Is there a correlation between the tRF fragment abundance (a majority originate from tRNAs with anticodons for G-ending codons) and translation efficiency in amoeba? The abundance of tRFs originating from different tRNAs varied during the development stages (analysis of *E. invadens* small RNA data Fig. 3C)-tRFs from tRNA^Gln_CTG^ were the most abundant tRFs in trophozoites whereas tRNA^Asp_GTC^ was the most abundant for late cysts. Further work would be required to understand if this tRF abundance is a cause or effect of the gene expression changes during stage conversion.

A majority of the tRFs identified through bioinformatic analysis mapped to the 5’ end of tRNAs. Similar results have been reported elsewhere such as in *Plasmodium falciparum* (14). One reason for the overrepresentation of 5’tRFs could be due to the nature of our library preparation-the libraries used in this analysis involved a 5’ phosphate dependent cloning step (these were not prepared for tRF analysis). If the cleavage of the tRNA yields fragments with a 5’-OH group, these fragments would not be efficiently cloned in the library. For example, mature tRNAs cleaved in the middle, will yield two fragments: the fragment mapping to the 5’ end will have the 5’ phosphate group of the original tRNA. However, the other fragment (which would map to the 3’ end) will have a 5’-OH group. Hence, the reads originating from the 5’ end would be preferentially selected in our datasets. Northern blot analysis confirmed that both 5’ and 3’ tRNA halves were the most abundant tRFs observed during stress. For most tRNAs (e.g. tRNA^Thr_TGT^, tRNA^Asn_GTT^, tRNA^Gly_GCC^ etc) both 5’ and 3’ tRNA halves were confirmed by northern blotting. However, for tRNA^Ala_CGC^ which was one of the most abundant tRFs in the sequence analyses, only the 5’ tRNA half was observed Fig. 2B. We found no tRNA where only the 3’ tRNA half was observed in the northern blotting. Additional issues in our bioinformatic analysis could arise since we have included all reads that map to our selected tRNA sequences. A more thorough approach would omit sequence reads that also map elsewhere in the genome (47). However, in the present study, our aim was largely to check for the presence of tRNA fragments from existing sequencing data, rather than quantifying and comparing the tRNA fragments between samples.

*Entamoeba* are exposed to a variety of stresses during their life cycle inside the human host. The anaerobic parasite can survive extreme conditions such as high oxygen content during tissue invasion, reactive oxygen and nitrogen species released by the host immune response, and fluctuations in glucose availability (48, 49). The parasite employs a variety of strategies to cope with these stresses including the regulation of antioxidant protein expression (such as thioredoxin) (50), or H_2_O_2_ responsive proteins (49, 51). Recently tRFs have been discovered to be important regulators of gene expression, particularly during stress.

tRNA halves were the most abundant species of tRFs that accumulated during stress in amoeba. Cleavage of tRNAs at the anticodon loop to generate tRNA halves has been extensively reported in other organisms and parasites. In *G. lamblia*, 46nt-long si-tRNAs were reported to accumulate during encystation. Both si-tRNAs and slightly shorter tRNA halves (∼36nt in length) were observed in response to nutritional stress (13). Importantly, the si-tRNA were representatives of the entire tRNA family and not restricted to a few members. We observed a similar accumulation of tRNA halves in response to both encystation and various stress inducers (oxidative stress, serum starvation, or heat shock). As in *Giardia*, almost the entire tRNA family seems to accumulate in response to stress. However, unlike *G. lamblia*, the cleavage of tRNAs during stress or during encystation were identical in amoeba, yielding the same tRNA halves during both events. Furthermore, we observed both 5’ and 3’ halves in amoeba with a few tRNA halves (tRNA^Ala_AGC^, tRNA^Ala_TGC^, tRNA^Asp_GTC^, and tRNA^Arg_TCT^ accounting for around 90% of all sequence reads). This result was similar to what was observed in *P. falciparum* where approximately 90% of the total tRFs were derived from tRNAs coding for Pro, Phe, Asn, Gly, Cys, Gln, His, and Ala(14).

We had previously reported that extracellular vesicles secreted from encysting amoeba can modulate the encystation rates in recipient cells undergoing encystation (3). In our present work, we found that amoebic EVs contained tRNA halves and that tRNA half constituted 50% of reads mapping to tRNAs in EV small RNA dataset. Bioinformatics analyses of EVs and cellular RNAs showed no major difference in the tRNA half profiles between the two samples and it appears that the composition of tRNA halves in the cells is very similar to the cargo of secreted EVs. Further work would be required to assess if the tRNA halves in EVs are important for regulating encystation in the recipient cells. The RNA cargo of extracellular vesicles secreted by *Trichomonas vaginalis* was very well characterized and found to contain predominantly 5’ tRNA halves. Moreover, the RNA content was seen to be internalized by the human host cells by lipid-raft mediated endocytosis, suggesting that tRNA half carried by parasitic EVs could be a possible mode of extracellular gene-modulation (52). Similarly, in both old and new world *Leishmania*, tRFs are highly enriched in EVs and point to a conserved packaging of tRFs in EVs (53).

Amoebic parasites have been known to have a functional RNAi pathway mediated through a population of 27nt antisense RNAs (25, 40, 54, 55), including 3 Ago proteins which have been shown to bind sRNAs (26). Our data show that tRFs are also associated with the Ago proteins in amoeba. Further studies would be required to assess if the association of tRFs with the Ago proteins is an additional mode of gene silencing in amoeba. Kumar et. al. carried out a comprehensive analysis of Ago PAR-CLIP data and showed that 5’ and 3’-tRFs can associate with target mRNAs in a mechanism reminiscent of microRNA gene silencing (37). With the bioinformatics analyses of our Ago-IP sequencing datasets, we also observed short reads (around 24-30nts in size) mapping to the tRNAs. The possibility of tRNA-derived fragments interacting with the RNAi pathways in amoeba offers an intriguing new role for the abundant tRNA genes in the amoebic genome.

Our data shows that various tRNA fragments are present in amoeba and that, similar to other organisms, accumulation of tRNA halves during stress-conditions is a mechanism that is conserved in amoeba. However, further work is required to figure out how exactly the tRNA halves accumulated during stress modulate protein synthesis and what the physiological role of this modulation is during amoebic infection.

## Materials and methods

### Parasite culture and cell lines

*E. histolytica* trophozoites (HM-1:IMSS) were grown axenically in TYI-S-33 (Trypticase, yeast extract, iron, and serum) medium in standard conditions described previously (56). *E. invadens* strain IP-1 was cultured in LYI-S-2 at 25°C (35). For encystation experiments, trophozoites at mid-log phase were iced, pooled, washed, and seeded into tubes in encystation medium (47% LYI-LG), and incubated at 25 °C. To measure encystation efficiency, total cells were counted using a hemocytometer before and after treatment with 0.1% sarkosyl. Parasites were transfected using the Attractene transfection reagent (Qiagen) to generate stable *E. histolytica* cell lines. All parasite-transfected lines were maintained at 6 μg/ml G418.

### Extracellular vesicle isolation

Parasites, grown to confluence in T25 flasks, were washed with serum-free TYI medium and incubated with 10 ml serum-free TYI medium for 16 hrs in an anaerobic chamber (BD GasPak EZ gas-generating container systems with GasPak EZ Campy container sachets; catalog number 260680). The conditioned medium was collected, centrifuged at 2,000 rpm, to remove cell debris and EVs pelleted using Total Exosome Isolation reagent (Invitrogen, Carlsbad CA, USA; catalog number 4478359) using manufacturer’s protocol. The EV pellet was further purified using size-exclusion chromatography with SmartSEC™ Mini EV Isolation System, (Systems Biosciences, CA) (3).

### RNA isolation and analysis

Cellular RNA was isolated using standard TRIzol (Invitrogen)-based protocol according to the manufacturer’s protocol. For isolation of RNA from Immunoprecipitation samples, 300 µl of TRIzol (Invitrogen) reagent was added to the final IP beads, and total RNA was isolated using manufacturer’s protocol.

### Immunoprecipitation

Immunoprecipitation experiments were done as previously described (28). Briefly, cell lysate was diluted with lysis buffer for a final protein concentration of 1 μg/μl. 20-30 μl packed anti-Myc beads (Thermo Scientific) were added to the IP mixture and rotated for 2 hours at 4 °C. The beads were washed 6 times at 4 °C (5 min each) using a low stringency wash buffer (the basic lysis buffer plus 0.1% (v/v) Tween-20, 0.1% (v/v) NP-40, 1 mM PMSF and 0.5X HALT EDTA free protease inhibitors). After the final wash step, the beads were pelleted and used for RNA preparation using TRIzol reagent.

### Northern blot analysis

Northern blot protocol was performed as previously described (25). 20 μg RNA samples were separated on a denaturing 12% polyacrylamide gel and transferred to a membrane (Amersham™ Hybond™ -N+ Membrane, GE Healthcare). Probe DNA was 5′-end labeled by PNK reaction using γ-[^32^P]-ATP and hybridized with the membrane in perfectHyb buffer (Sigma) overnight at 37 °C. The membrane was washed using low stringency condition (2X SSC, 0.1% SDS at 37 °C for 20 min) and medium stringency condition (1X SSC, 0.1% SDS at 37 °C for 20 min). Radioactive signal was detected using a Phosphor screen and imaged on a Personal Molecular Imager (Bio-Rad). The various probes used for corresponding tRFs are in the following table:

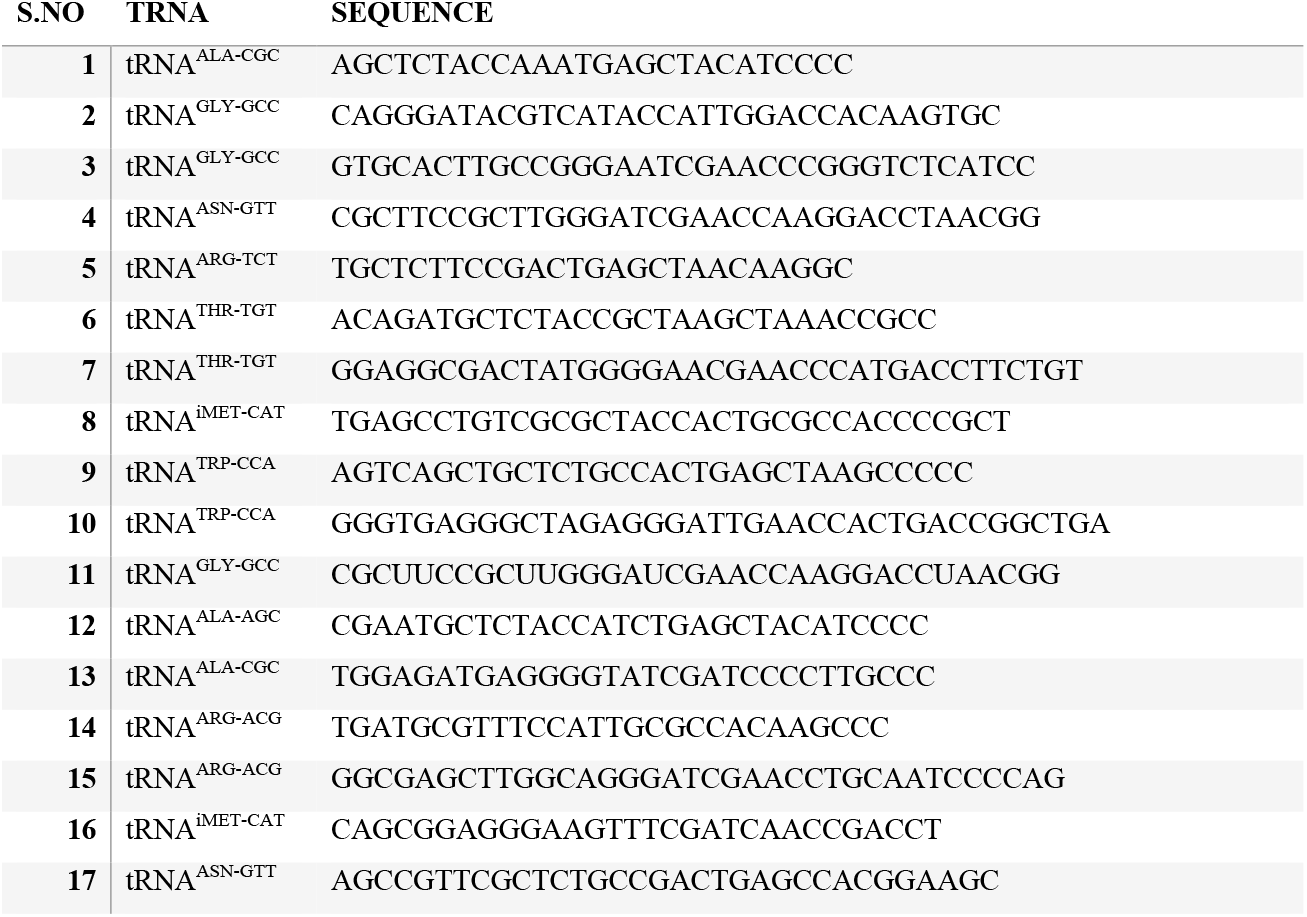

### Bioinformatics analyses

RNA sequencing data were analyzed using a previously described pipeline (40). Raw reads were processed using cutadapt (57). Sequences were then mapped to *E. histolytica* tRNA using Bowtie v1.2.2 (http://bowtie-bio.sourceforge.net) with the parameters −v1 and −best (30). tRNAscan-SE was used to predict tRNA genes from the genomes of *E. histolytica* and *E. invadens* (29). CCA was added to the genes at the 3’ end.

tRNAscan-SE command used in this prediction run:

tRNAscan-SE -qQ --detail -o# -m# -f# -l# -c <u>tRNAscan-SE.conf</u> -b# -s# (fasta file)

## Acknowledgements

The project was supported by funds to U.S. from the National Institutes of Health (R01-AI121084 and R21-AI125764). We thank all members of the Singh lab for their help and contribution to this work. Its contents are solely the responsibility of the authors and do not necessarily represent the official views of the National Institutes of Health.

## Supplementary Figures

Fig. S1: Northern blot analysis of total RNA from 24 h cysts and trophozoites probed for tRNA halves. tRNA halves were seen to accumulate for the different tRNAs probed.

Fig. S2: IGV mapping of reads mapping against tRNA templates-tRNA^Ala_TGC^ and tRNA^Arg_TCT^. A 24nt 5’ tRF is seen to be associated with all Ago proteins for tRNA^Ala_TGC^. An itRF species is seen to be associated with Ago2-2 for both tRNAs.

